# Activated STING in the thymus alters T cell development and selection leading to autoimmunity

**DOI:** 10.1101/2024.02.17.580803

**Authors:** Zimu Deng, Christopher S. Law, Santosh Kurra, Noa Simchoni, Anthony K. Shum

## Abstract

Classifying systemic inflammatory disorders as autoinflammatory or autoimmune provides insight into disease pathogenesis and whether treatment should target innate molecules and their signaling pathways or the adaptive immune response. COPA syndrome is a monogenic disorder of immune dysregulation that leads to interstitial lung disease and high-titer autoantibodies. Studies show constitutive activation of the innate immune molecule STING is centrally involved in disease. However, the mechanisms by which STING results in loss of T cell tolerance and autoimmunity in COPA syndrome or more common autoimmune diseases is not understood. Using *Copa^E241K/+^* mice, we uncovered a functional role for STING in the thymus. Single cell data of human thymus demonstrates STING is highly expressed in medullary thymic epithelial cells (mTECs) involved in processing and presenting self-antigens to thymocytes. In *Copa^E241K/+^* mice, activated STING in mTECs triggered interferon signaling, impaired macroautophagy and caused a defect in negative selection of T cells. Wild-type mice given a systemic STING agonist phenocopied the selection defect and showed enhanced thymic escape of a T cell clone targeting a self-antigen also expressed in melanoma. Our work demonstrates STING activation in TECs shapes the T cell repertoire and contributes to autoimmunity, findings important for settings that activate thymic STING.

**Summary:** Activation of STING in thymic epithelial cells shifts the T cell repertoire to promote autoimmunity providing a new mechanistic link between innate and adaptive immunity.

## Introduction

The use of genetic sequencing to study monogenic inborn errors of immunity (IEI) has allowed researchers to pinpoint specific molecular pathways defective in disease. These insights have enabled the classification of immunological disorders into autoimmune and autoinflammatory diseases, depending on whether the adaptive or innate arm of the immune system is dysregulated, respectively. In practice, these disorders, as well as more common conditions with complex genetics such as rheumatoid arthritis or systemic lupus erythematosus, exist on a spectrum between autoimmunity and autoinflammation and many share features of both. Further investigation is needed to unravel the specific mechanisms by which overactivation of the innate immune system in autoinflammatory conditions reprograms the adaptive response to drive autoimmune disease (1).

COPA syndrome is a monogenic disorder of immune dysregulation that presents with interstitial lung disease and high-titer autoantibodies (2). Patients may also have pulmonary hemorrhage, renal disease and inflammation of small and large joints. Treatment of COPA syndrome involves high dose immunosuppression which can decrease episodes of life-threatening alveolar hemorrhage and severity of other clinical manifestations (3), although some patients develop end-stage pulmonary fibrosis that is refractory to therapy and requires lung transplantation (4).

To define the mechanisms of COPA syndrome, we recently established a mouse model by generating *Copa^E241K/+^* knock-in mice that express one of the same *Copa* missense mutations as patients (5). *Copa^E241K/+^* mice spontaneously developed clinical and immunologic features of patients, including interstitial lung disease and increased levels of activated, cytokine-secreting T cells. Bone marrow chimera and thymic transplant experiments showed that expression of mutant COPA in the thymic stroma was sufficient to cause a defect in negative selection of CD4^+^ T cells. Adoptive transfer of T cells harvested from *Copa^E241K/+^* mice into immunodeficient animals caused organ autoimmunity. Our data suggested that a key step in the initiation of disease in COPA syndrome is a breakdown in central tolerance. However, the mechanisms by which mutant COPA might lead to impaired thymic tolerance remain unknown (5).

COPA is one of seven subunits of coat protein complex I (COPI) that mediates retrograde movement of proteins from the Golgi apparatus to the endoplasmic reticulum (ER). Pathogenic missense mutations that disrupt the WD40 domain of COPA impair the binding and sorting of dilysine-tagged client proteins targeted for retrieval to the ER in COPI vesicles (2). We and others recently found that mutant COPA causes mis-trafficking of the innate immune adapter molecule STING. STING is a signaling receptor protein in the cytosolic DNA sensing pathway of cells (6–8). STING is activated by the second messenger cyclic GAMP (cGAMP), a ligand generated by the enzyme cyclic GMP-AMP synthase (cGAS) after it binds to double stranded DNA. Upon binding cGAMP or other cyclic dinucleotides, STING traffics to the ER-Golgi intermediate compartment (ERGIC) and Golgi where it activates the kinase TBK1 to induce type I interferons and other inflammatory cytokines. In COPA syndrome, defective retrieval of STING by mutant COPA results in the accumulation of STING on the Golgi and persistent STING signaling (6, 9, 10).

Although aberrant STING activation has been linked to several autoimmune diseases including more common disorders such as rheumatoid arthritis and systemic lupus erythematosus (SLE) (11, 12), the specific mechanisms by which STING leads to a loss of self-tolerance and autoreactive T cell responses is not well understood. Similar to SLE, peripheral blood mononuclear cells from COPA syndrome subjects exhibit a markedly elevated type I interferon stimulated gene (ISG) signature (13). *Copa^E241K/+^* mice phenocopy the type I interferon (IFN) activation observed in patients, not only in immune cells, but also in the thymic epithelium (6). Remarkably, all the abnormal T cell phenotypes and interferon signaling are completely reversed after crossing *Copa^E241K/+^* mice to STING-deficient *Goldenticket* mice (*Sting1^gt/gt^*). Taken together, our data suggests STING may have a functional role in the thymus that predisposes to autoimmunity in COPA syndrome. In this study, we hypothesized that activation of STING in the thymic epithelium leads to a break in self-tolerance by directly altering T cell development and selection.

## Results

### Activated STING in thymic stroma of Copa^E241K/+^ mice upregulates interferons

To assess whether STING has a functional role in the thymic stroma of *Copa^E241K/+^* mice that influences the development and selection of autoreactive T cells, we first sought to determine whether we could detect activation of STING protein in TECs. We performed western blots of enriched TEC cell populations and assayed for phosphorylated STING (pSTING). We found a significant increase in pSTING in TECs of *Copa^E241K/+^* mice in comparison to wild-type mice (**Fig. 1A**) and used confocal microscopy to confirm that mTECs in *Copa^E241K/+^* mice had evidence of pSTING (**Fig. 1B**). Consistent with these results, we had shown previously that bulk RNA sequencing of sorted TEC populations from *Copa^E241K/+^* mice have an increase in *ifnb1* transcript levels, particularly in MHC-II^high^CD80^high^ medullary thymic epithelial cells (mTEC^high^) (6). We performed deeper analysis of mTEC^high^ cell bulk RNA sequencing data to determine whether other signaling outputs of STING were upregulated. Interestingly, although *ifnb1* transcript was elevated in *Copa^E241K/+^* mice, the cytokines *il6* and *tnf* that are downstream of NF-κB signaling were not (**Fig. 1C**). Because activated STING is also associated with cellular senescence (14) and apoptosis (15), we examined tissue architecture of thymi by confocal microscopy to assess whether there was evidence of thymic involution. Immunofluorescence staining with KRT5 and KRT8 cell surface markers revealed that *Copa^E241K/+^* mice appeared to have normal size and organization of the cortical and medullary thymic epithelium (16) (**Fig. 1D**). Furthermore, the thymi from *Copa^E241K/+^* mice had normal medullary and cortical thymic epithelial cell (mTEC and cTEC) numbers, suggesting an absence of an increase in cell death (**Fig. 1E**). Thus, constitutive activation of STING in thymic epithelial cells appears to cause upregulation of IFNs but other STING-associated signaling outputs including NF-κB activation and cell death did not appear to be affected. These data align with prior work showing that there are context (17) and cell-type specific effects of STING that are partially influenced by signal strength (18).

**Figure 1.**
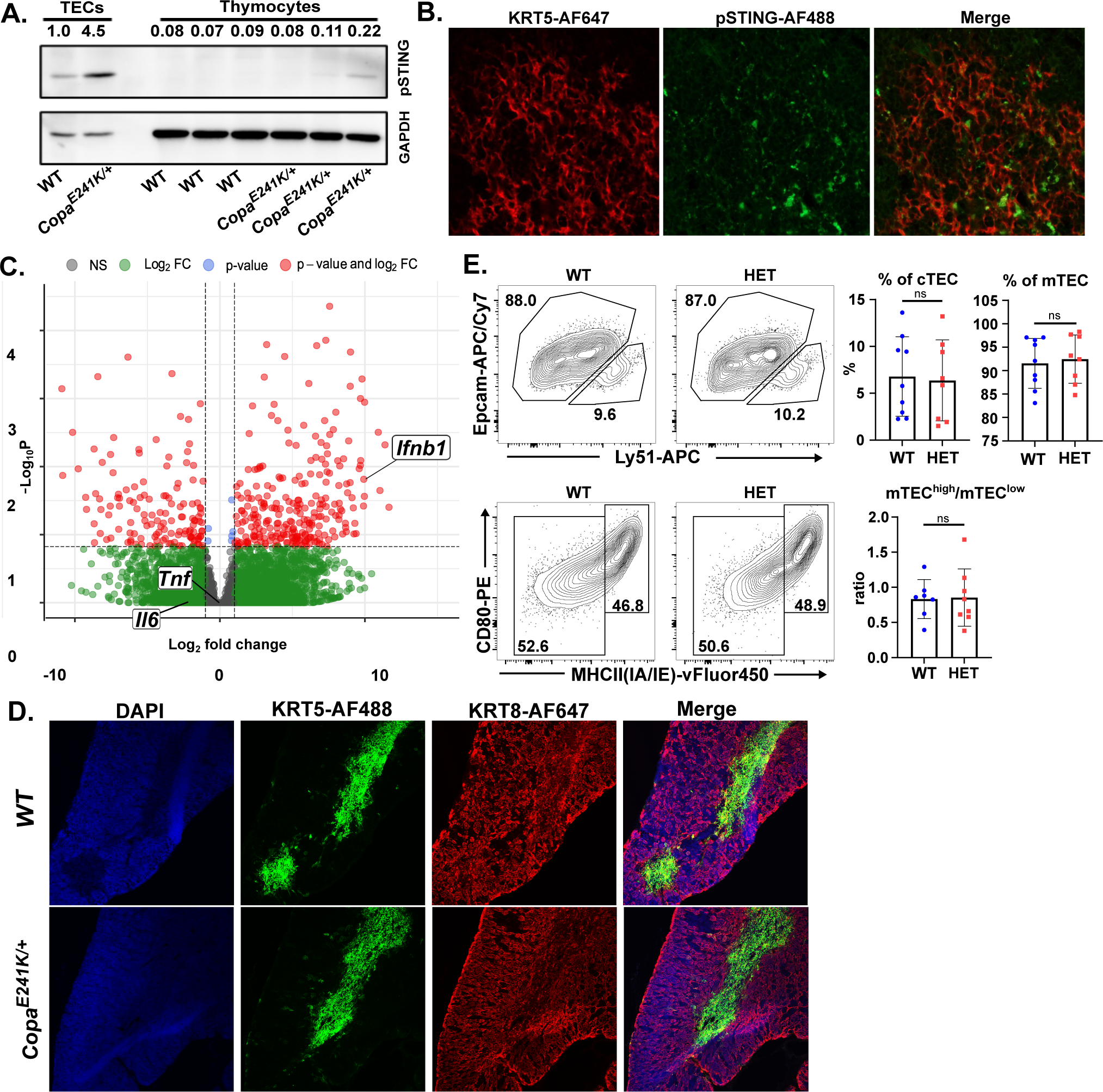
Activated STING in thymic stroma upregulates interferon signaling. **(A)** Representative immunoblot of phosphorylated STING (pSTING) in thymic epithelial cells (TECs) versus thymocytes with relative band density quantification. Data represents 2 independent experiments. **(B)** Immunofluorescence stain of keratin 5 (KRT5) and pSTING expression on thymic sections from *Copa^E241K/+^* mice. **(C)** Volcano plot of RNA sequencing analysis of sorted medullary TECs (mTECs) from WT and *Copa^E241K/+^* mice (n = 3 per genotype). **(D)** Immunofluorescence stain of KRT5 and keratin 8 (KRT8) on thymic sections from *Copa^E241K/+^* and WT littermates. **(E)** Top left: flow cytometry of Ly51 vs EpCAM on TECs; top right: percentages of mTECs and cortical TECs (cTECs); bottom left: mTEC expression of CD80 and MHC-II; bottom right: ratio of mTEC^high^ versus mTEC^low^. Data are mean ± SD. Unpaired, parametric, two-tailed Student’s t-test was used for statistical analysis. p < 0.05 is considered statistically significant. ns: not significant.

### Activated STING in thymic stroma increases post-selection thymocytes

Having found that STING is activated in the thymic epithelium of *Copa^E241K/+^* mice, we next wanted to assess how STING influenced thymocyte development. As in our previous study (5), we observed that *Copa^E241K/+^* mice have an increase in single positive (SP) thymocytes (**Supplementary Fig. 1A**) and post-selection thymocytes (**Supplementary Fig. 1B**). To assess whether activated STING triggered these changes, we examined STING-deficient *Copa^E241K/+^* mice (*Copa^E241/+^×Sting1^gt/gt^* mice) and found that the increase in the thymocyte populations was completely reversed (**Supplementary Fig. 1A-B**). We next used bone marrow chimeras to determine whether activated STING in the radioresistant stromal compartment or the hematopoietic compartment caused the changes to the thymocyte populations, recognizing that STING is expressed not only in T cells, but also other thymic antigen presenting cells involved in selection (19, 20). We transplanted wild-type bone marrow into irradiated *Copa^E241K/+^* mice and found an increase in SP (**Fig. 2A**) and post-selection thymocytes (**Fig. 2B**) similar to unmanipulated *Copa^E241K/+^* mice. In contrast, the alterations to the thymocyte populations were reversed when wild-type bone marrow was transplanted into irradiated *Copa^E241/+^×Sting1^gt/gt^* mice (**Fig**. **2A-B**), suggesting that STING exerts its influence on thymocyte populations from within the thymic stroma.

**Figure 2.**
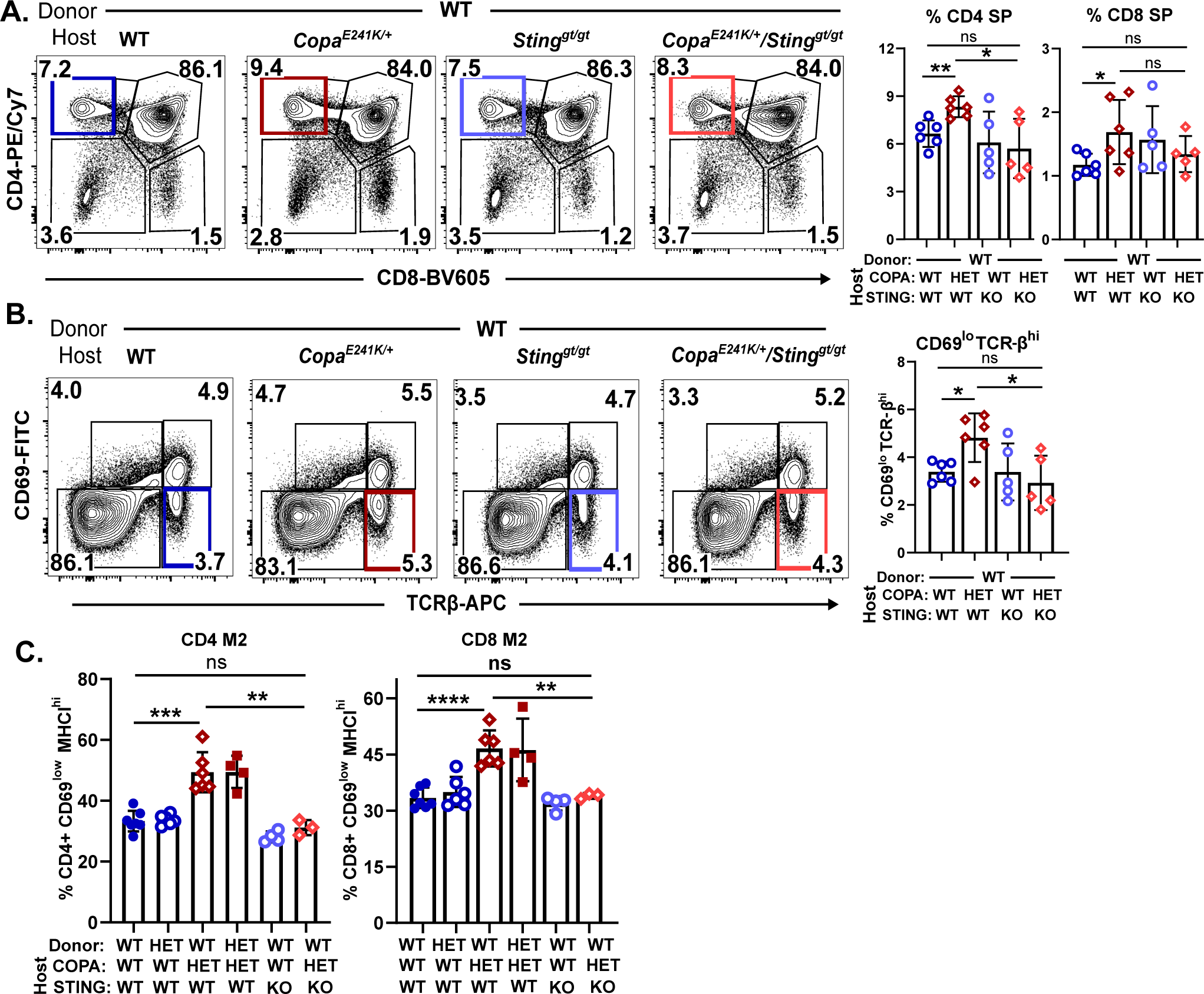
Activated STING in thymic stroma increases single positive thymocytes. **(A)** Left: representative flow plots of CD4 and CD8 on reconstituted thymocytes in bone marrow chimeras. Right: percentages of CD4 SP and CD8 SP thymocytes among the reconstituted thymocytes (*WT→WT*, n = 6; *WT→Copa*^E241K/+^, n = 6; *WT→Sting*^gt/gt^, n = 5; *WT→Copa*^E241K/+^ *× Sting*^gt/gt^, n = 5). **(B)** Left: representative flow analysis of CD69 and TCRβ on reconstituted thymocytes in bone marrow chimeras. Right: percentages of CD69^high^TCRβ^high^ and CD69^low^TCRβ^high^ among the reconstituted thymocytes **(C)** Percentages of CD69^low^MHCI^high^ (mature stage 2: M2) among the reconstituted CD4 and CD8 single positive thymocytes. Flow strategy in Supplementary Fig. 1D. (*WT→WT*, n = 7; *WT→Copa*^E241K/+^, n = 6; *Copa*^E241K/+^*→WT*, n = 6; *Copa*^E241K/+^*→Copa*^E241K/+^, n = 4; *WT→Sting*^gt/gt^, n = 4; *WT→Copa*^E241K/+^ *× Sting*^gt/gt^, n = 3). Data are mean ± SD. Unpaired, parametric, two-tailed Student’s t-test was used for statistical analysis. p < 0.05 is considered statistically significant. ns: not significant. HET: *Copa^E241^*^K/+^, Sting KO: *Sting*^gt/gt.^

Having identified a role for STING in TECs, we next wanted to examine how its effect on interferon signaling influenced thymocyte maturation. Our prior work suggested that higher type I IFN levels in thymic epithelial cells of *Copa^E241K/+^* mice imprints thymocytes with elevated interferon stimulated genes (ISGs) and promotes an expansion of late-stage SP cells (6). To assess whether thymic epithelial STING mediates these changes in *Copa^E241K/+^* mice, we again examined bone marrow chimeras for reconstituted thymocyte populations. First, we found that STING in thymic stromal cells of *Copa^E241K/+^*mice (**Supplementary Fig. 1C**) upregulated ISGs in thymocytes. We then used a flow cytometry staining strategy that subsets increasingly mature SP cells into semi-mature (SM), mature 1 (M1), and mature 2 (M2) populations (21) (**Supplementary Fig. 1D**) Consistent with the results from above, we found that loss of STING in the thymic stroma was able to reverse the expansion of M2 cells in the *Copa^E241K/+^* thymus (**Fig. 2C, Supplementary Fig. 1E**). Taken together, these data establish that chronic STING signaling in thymic epithelial cells results in an increase in post-selection SP thymocytes and promotes the expansion of late-stage SP thymocytes.

To dissect out how STING-mediated type I IFN signaling altered thymocyte development, we next performed bone marrow chimeras using interferon receptor-deficient mice (*Ifnar^-/-^*). We transferred bone marrow from *Ifnar^-/-^* mice into irradiated wild-type or *Copa^E241K/+^* hosts. Loss of IFNAR on thymocytes completely blunted the upregulation of ISGs caused by the *Copa^E241K/+^* thymic stroma (**Supplementary Fig. 2A**). However, the expansion of post-selection SP thymocytes was not fully rescued by IFNAR deficiency, suggesting that chronic STING signaling altered thymocyte development through additional mechanisms (**Supplementary Fig. 2B-C**). Although a growing body of work indicates transient STING activation induces non-canonical autophagy (22, 23), the role of STING on autophagic function in TECs has not been explored. Based on our observations in prior work (2, 5), we speculated that chronic STING activation in TECs might actually impair macroautophagy (hereafter autophagy), a cellular process critical for processing and presenting self-peptides to thymocytes, particularly during negative selection (24). We reasoned that if persistent STING activation in *Copa^E241K/+^* mice did indeed disrupt autophagic function in TECs, this might account for the increase in post-selection SP thymocytes that persisted despite loss of IFNAR on thymocytes.

### Chronic STING signaling in TECs impairs autophagy

To investigate the role of chronic STING activation on autophagy in our model, we crossed *Copa^E241K/+^* mice to GFP-LC3 reporter mice (25) and measured GFP-LC3II^+^ by flow cytometry in TECs. Interestingly, mTEC^high^ cells in *Copa^E241K/+^*mice had a striking increase in the percentage of cells expressing autophagosome-associated GFP-LC3II (**Fig. 3A**). To determine whether the increase in autophagosomes reflected enhanced autophagy initiation or impaired autophagic flux, we expressed wild-type and E241K mutant COPA in HEK293T cells that stably express STING. We cultured the cells with and without bafilomycin A (BafA) which inhibits autophagosome/lysosome fusion (26). In cells expressing wild-type COPA we observed an expected increase in LC3II levels after treatment with BafA. In cells expressing E241K mutant COPA, however, we observed higher levels of LC3II at baseline that did not increase in response to BafA treatment, consistent with impaired autophagic flux (**Fig. 3B**).

**Figure 3.**
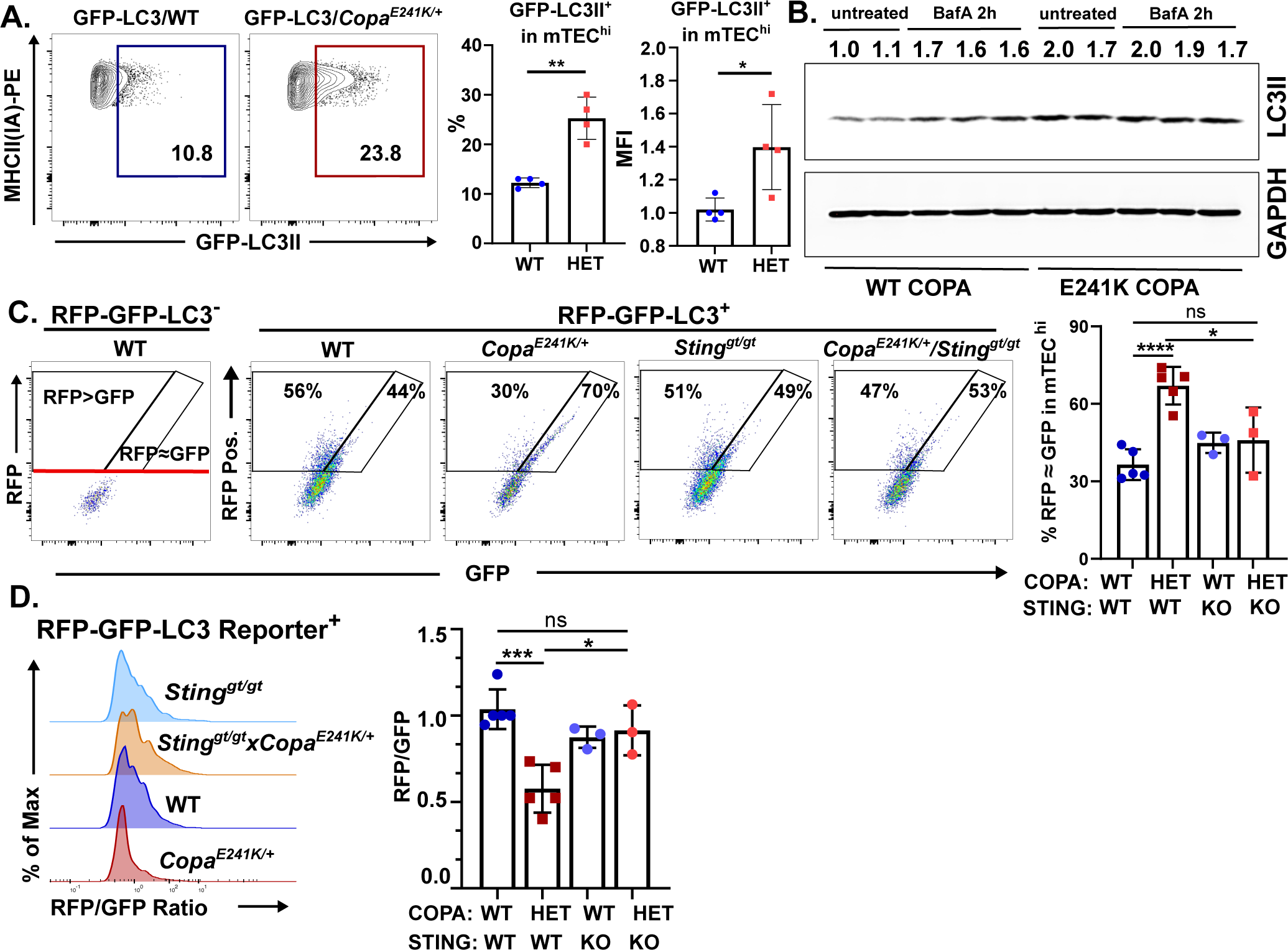
Constitutive activation of STING in the thymus impairs autophagic flux. **(A)** Left: Flow analysis of autophagosome-associated LC3 (LC3II) in mTEC^high^ from GFP-LC3 and GFP-LC3 × *Copa^E241K/+^* mice. Right: percentage of LC3II-GFP+ population among total mTEC^high^ and GFP MFI in mTEC^high^. (GFP-LC3 × WT, n = 4, GFP-LC3 × *Copa^E241K/+^*, n = 4). **(B)** Representative immunoblot and densitometric analysis of LC3II following transient transfection of WT or E241K COPA in HEK293T cells that stably express STING. **(C)** Quantitation of autophagic flux in mTECs of *CAG-RFP-GFP-LC3* tandem reporter mice. Left: flow cytometry of autophagosome-(RFP≈GFP) and autolysosome-associated (GFP<RFP) LC3 in mTECs. Right: percentage of mTEC^high^ with reduced autophagic flux (RFP-GFP-LC3 × WT, n = 5, RFP-GFP-LC3 × *Copa^E241K/+^*, n = 5; RFP-GFP-LC3 × WT × *Sting^gt/gt^*, n = 3, RFP-GFP-LC3 × *Copa^E241K/+^ × Sting^gt/gt^*, n = 3). (D) Left: ratio of RFP/GFP fluorescence histogram in mTECs expressing LC3 tandem reporter; and right: mean RFP/GFP ratio in mTECs (RFP-GFP-LC3 × WT, n = 5, RFP-GFP-LC3 × *Copa^E241K/+^*, n = 5; RFP-GFP-LC3 × *WT × Sting^gt/gt^*, n = 3, RFP-GFP-LC3 × *Copa^E241K/+^* × *Sting^gt/gt^*, n = 3). Data are mean ± SD. Unpaired, parametric, two-tailed Student’s t-test was used for statistical analysis. p < 0.05 is considered statistically significant. ns: not significant HET: *Copa^E241K/+^*, Sting KO: *Sting^gt/gt^*.

To evaluate autophagic function *in vivo*, we crossed *Copa^E241/+^*mice with *CAG-RFP-EGFP-LC3* tandem reporter mice so that we could quantitate autophagic flux in TECs by flow cytometry (27, 28). Because EGFP is more sensitive than RFP to the acidic environment of autolysosomes, cells with higher flux have less EGFP and thus an increased RFP/GFP ratio (**Fig. 3C-D**). Consistent with our data above, *Copa^E241/+^* mTEC^high^ cells had a lower RFP/GFP ratio than wild-type mTEC^high^ cells, indicating impaired autophagic flux. In addition, the RFP/GFP ratio returned to wildtype levels in STING-deficient *Copa^E241/+^/Sting1^gt/gt^* mice (**Fig. 3C-D**). Collectively, these findings indicate that chronic STING activation impairs autophagic flux which may in turn alter the delivery of endogenous self-peptides to developing T cells.

### Chronic STING activation in TECs impairs T cell selection and alters the T cell repertoire

Having found that chronic STING activation impaired autophagic flux in mTECs of *Copa^E241/+^* mice, we next wanted to examine the impact of this on repertoire selection of T cells in our model. Autophagy in mTECs facilitates the loading of self-peptides onto MHC class II molecules for presentation to thymocytes auditioning for selection (20, 24). To study this, we employed the RiP-mOVA/OT-II system to study negative selection of T cells. RiP-mOVA mice express membrane bound ovalbumin (mOVA) as a neo-self-antigen within mTECs and OT-II cells have a T cell receptor (TCR) specific for ovalbumin peptide. Normally, when OT-II cells traffic through the thymus and encounter ova peptide presented by MHC-II on mTECs, they undergo clonal deletion and die (29).

We transplanted bone marrow from OT-II mice into RiP-mOVA mice bred onto the *Copa^E241K/+^*, *Copa^+/+^/Sting1^gt/gt^*and *Copa^E241K/+^/Sting1^gt/gt^* genetic backgrounds and analyzed clonal deletion of OT-II thymocytes. In *Copa^E241K/+^* mice, we found a defect in negative selection of CD4 T cells compared to wild-type mice, characterized by an increase OT-II CD4 SP thymocytes (**Fig. 4A-B**). Remarkably, loss of STING in *Copa^E241K/+^/Sting1^gt/gt^* mice restored the percentages of CD4SPs to levels comparable to wild-type mice (**Fig. 4A-B**). We evaluated clonotypic CD4 SP cells for expression of cell surface CD5 and intracellular Nur77, an immediate early gene rapidly up-regulated by TCR signaling (30). In OT-II cells from *Copa^E241/+^*mice, we found that CD5 and Nur77 were significantly lower than in both wild-type and *Copa^E241K/+^/Sting1^gt/gt^* mice (**Fig. 4C, Supplementary Fig. 3A**). Because Vβ5 and Vα2 expression remained intact (**Fig. 4B**), this suggested the cells did not engage their TCR with peptide-MHC-II because impaired autophagy in mTECs disrupted the proper processing and presentation of peptide antigen, which overall lead to an increase in post-selection SP thymocytes that escaped negative selection.

**Figure 4.**
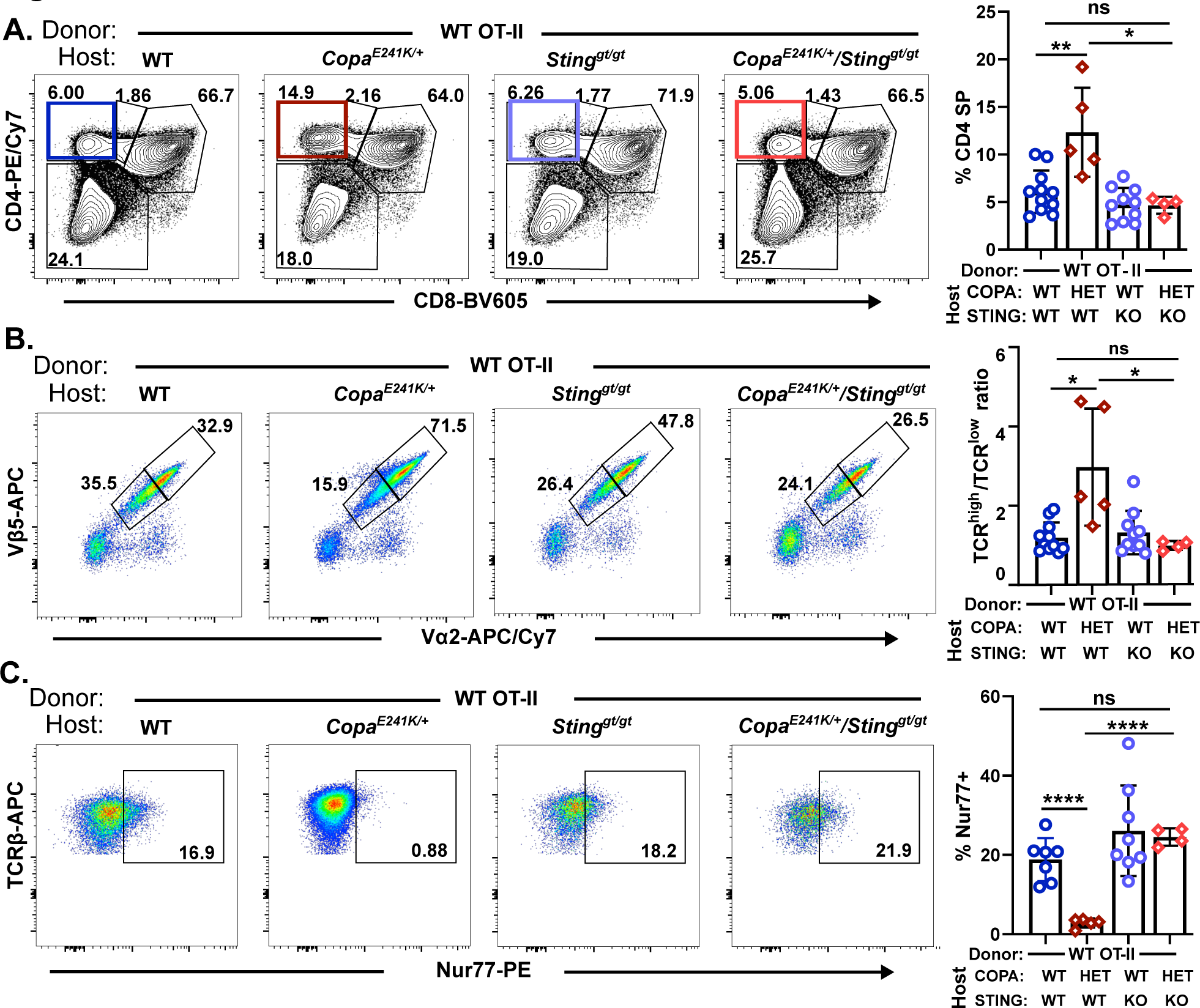
Activated STING impairs negative selection of T cells and alters the T cell repertoire. **(A)** Left: representative flow analysis of CD4 and CD8 on reconstituted thymocytes in bone marrow chimeras. Right: percentage of CD4 single positive thymocytes among the reconstituted thymocytes (OTII→Rip-mOVA × WT, n = 11; OTII→Rip-mOVA × *Copa^E241K/+^*, n = 5; WT→Rip-mOVA × WT *× Sting^gt/gt^*, n = 10; WT→Rip-mOVA × *Copa^E241K/+^ × Sting^gt/gt^*, n = 4). **(B)** Left: flow analysis of Vβ5 and Vα2 in CD4 SP thymocytes in the bone marrow chimeras shown in **(A)**. Right: ratio of TCR^high^ versus TCR^low^ among CD4 SP thymocytes. **(C)** Left: flow analysis of TCRβ and Nur77 expression in the reconstituted CD4 SP in the bone marrow chimeras. Right: percentage of Nur77+ population among CD4 SP thymocytes. (OTII→Rip-mOVA × WT, n = 7; OTII→Rip-mOVA *× Copa^E241K^*^/*+*^, n = 5; WT→Rip-mOVA × WT × *Sting^gt/gt^*, n = 8; WT→Rip-mOVA × *Copa^E241K/+^ × Sting^gt/gt^*, n = 4). Data are mean ± SD. Unpaired, parametric, two-tailed Student’s t-test was used for statistical analysis. p < 0.05 is considered statistically significant. ns: not significant. HET: *Copa^E241K/+^*, Sting KO: *Sting^gt/gt^*, all host mice are Rip-mOVA background.

To determine whether the defect in selection we observed was limited only to mOVA and OT-II cells or could be applied more broadly, we first assessed the CD5 level on polyclonal SP thymocytes and found SP thymocytes in *Copa^E241K/+^* mice have a modest but significant drop in CD5 on SP cells (**Supplementary Fig. 3B**), suggesting the selection of T cells across a broad range of specificities might be affected. Next, we compared TCRVβ chain profiles in WT and *Copa^E241K/+^* mice and found a shift in the TCRVβ repertoire of CD4 SP and CD8 SP thymocytes in *Copa^E241K/+^* mice, which were reversed in *Copa^E241K/+^/Sting1^gt/gt^* mice (**Supplementary Fig. 4C**). Taken together, our findings suggest that chronic STING activation causes a defect in negative selection of T cells and an overall shift in the immune cell repertoire.

### A systemic STING agonist increases autoreactive T cells in the thymus

Having found that constitutive STING activation in thymic epithelial cells alters thymocyte development and selection, we wondered if our findings might be applied more broadly outside the context of the COPA syndrome model. Specifically, we sought to examine whether activation of STING in the thymic stroma using a systemic STING agonist developed for clinical use (for instance, cancer immunotherapy) would phenocopy our findings and promote the output of autoreactive T cell clones. To study this, we turned to TRP-1 TCR Tg mice, an MHCII-restricted mouse model in which CD4^+^ T cells recognize an epitope of the endogenous tyrosinase-related protein 1 (TRP-1) (31). TRP-1 protein is expressed as a melanocyte self-antigen in medullary thymic epithelial cells and also in melanoma tumors. Expression of TRP-1 in mTECs results in the deletion of TRP-1 specific T cells in the thymus and prevents their escape into peripheral lymphoid organs (16, 32). Because our data suggested that activation of STING in mTECs leads to a defect in negative selection, we set out to study whether systemic delivery of a STING agonist would disrupt processing and presentation of the TRP-1 self-antigen to developing thymocytes. To assess this, we treated TRP-1 TCR Tg mice (on a *Rag^-/-^*background) with the recently reported diABZI STING agonist (33) every other day for 2 weeks (**Supplementary Fig. 4A**). After a brief washout period, we assessed the thymus for the selection of TRP-1 specific T cells. In mice treated with the diABZI STING agonist, we found a significant increase in the percentages of CD3^+^Vβ14^+^ T cells in comparison to control mice treated with vehicle alone (**Fig. 5A**). An examination of CD3^+^Vβ14^+^ signaled thymocytes showed significantly lower levels of cleaved caspase-3 in mice receiving the STING agonist, consistent with a decrease in clonal deletion of anti-tumor T cells in treated mice (34) (**Fig. 5B**).

**Figure 5.**
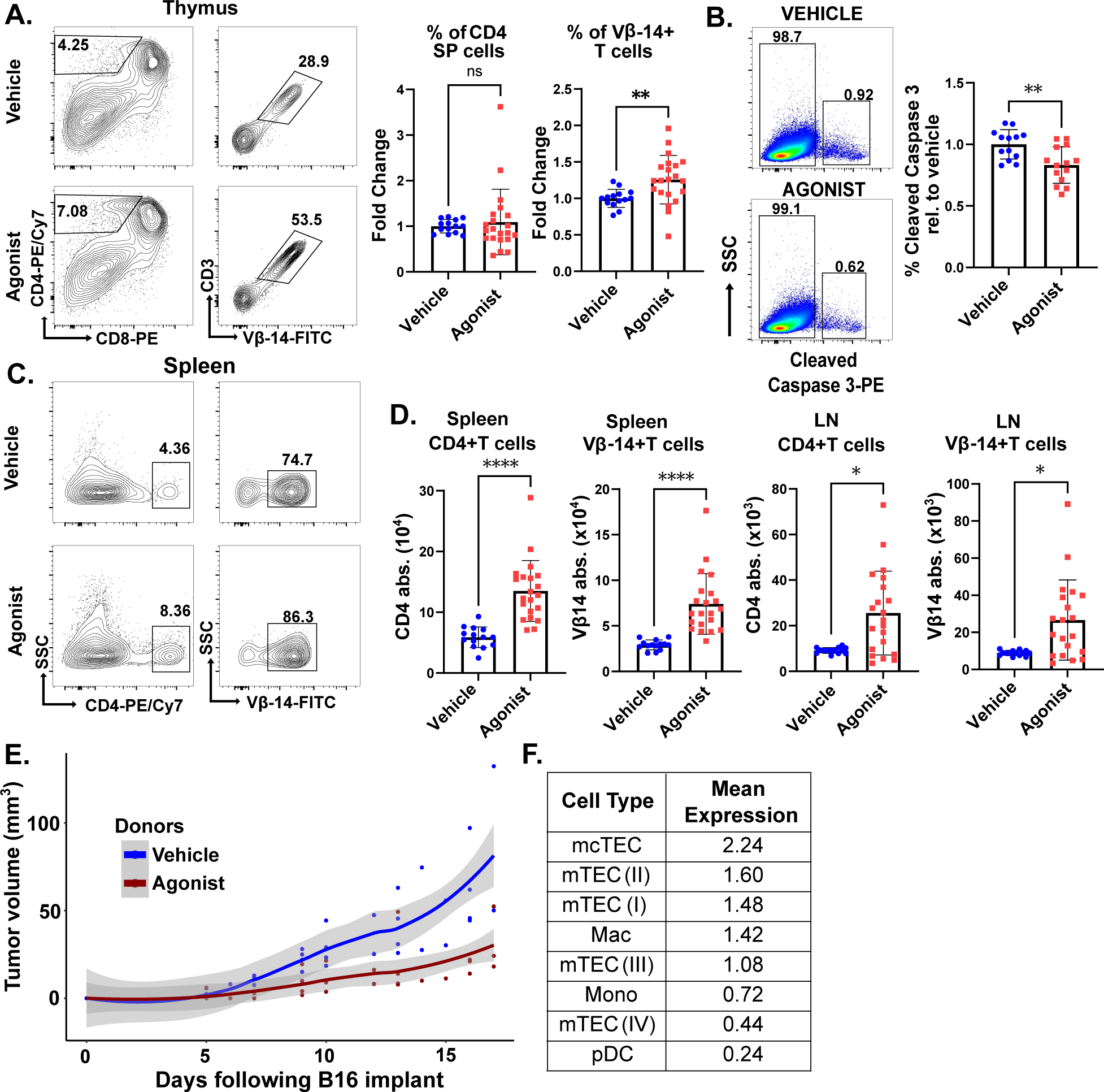
A systemic STING agonist increases autoreactive T cells in the thymus. **(A)** Left: CD4 and CD8 profile of thymocytes and CD3 and Vβ14 expression of CD4 single positive thymocytes in *Rag1^-/-^ Tyrp1^B-w/wt^* TCR mice treated with diABZI STING agonist or vehicle. Right: change in percentage of total thymic CD4 SP and Vβ14^+^ CD4 SP following STING agonist treatment (vehicle n = 14; agonist n = 21). **(B)** Left: cleaved caspase 3 on CD5^high^ TCRβ^high^ thymocytes in vehicle and agonist treated mice. Right: percentage of thymocytes undergoing clonal deletion in agonist treated mice relative to vehicle treatment (vehicle n = 13, agonist n = 14). Unpaired, parametric, two-tailed Student’s t-test was used for statistical analysis. **(C)** Left: flow analysis of splenic CD4 SP Vβ14^+^ autoreactive T cells in vehicle and agonist treated mice. Right: absolute number of CD4 SP Vβ14^+^ autoreactive T cells. **(D)** Absolute number of inguinal CD4 SP Vβ14^+^ autoreactive T cells in vehicle and agonist treated mice (vehicle n = 14, agonist n = 21). **(E)** B16 melanoma growth in *Rag1^-/-^*mice that received X-ray radiation, PD-1 antibody and adoptively transferred splenocytes from *Rag1^-/-^ Tyrp1^B-w/wt^* TCR mice treated with STING agonist or vehicle (vehicle n = 4, agonist n = 5). Locally estimated scatterplot smoothing with 95% confidence interval of B16 tumor growth over time. Data are mean ± SD. Two-tailed Mann-Whitney U-test was used for statistical analysis unless indicated above. p < 0.05 is considered statistically significant. ns: not significant. **(F)** Mean expression of *STING1* transcript in select cell types in human thymus.

We assessed peripheral lymphoid organs for the presence of CD3^+^Vβ14^+^ T cells. In lymph nodes and spleen, there was a significant increase in CD3^+^Vβ14^+^ T cells that escaped negative selection in the thymus (**Fig. 5C**). To confirm that these T cells were functionally competent we next sought to determine whether the cells were capable of mediating an anti-tumor immune response because although TRP-1 is a self-antigen, it is also expressed in B16 melanoma tumors. After the washout period, we isolated secondary lymphoid organs from TRP-1 TCR Tg mice treated with STING agonist or control vehicle to collect total CD3^+^Vβ14^+^ T cells. From each treated mouse, we adoptively transferred harvested cells into a *Rag1^-/-^* mouse implanted with B16 melanoma tumor. We observed significant regression of tumors in recipient mice that received CD3^+^Vβ14^+^ T cells from STING agonist treated mice in comparison to tumors in mice that received control cells (**Fig. 5E**). Taken together, activation of STING in the thymus independent of COPA syndrome leads to the escape of self-antigen specific T cells that otherwise normally undergo negative selection, findings relevant for other immunoregulatory disorders and in patients receiving small molecule drugs to modulate STING signaling during cancer immunotherapy.

### STING is highly expressed in medullary thymic epithelial cells in humans

Finally, although the *Copa^E241/+^* mouse model closely phenocopies COPA syndrome in patients and is a powerful model to study the disease (5, 6), we next wanted to directly determine the potential relevance of thymic STING in human tissue and assessed the levels of STING transcript in human thymus. We analyzed single cell RNA sequencing data from a human thymus cell atlas and found STING was variably expressed depending on the specific cell type (35). Among stromal cells, STING was most highly expressed in the thymic epithelium and exhibited significantly elevated expression in medullary thymic epithelial cells (mTECs), including those directly involved in processing and presenting tissue antigens to developing T cells (36) (**Figure 5F, Supplementary Figure 4C**). Interestingly, STING transcript levels in human mTECs was higher than that found in thymic macrophages and more than double the levels of monocytes or plasmacytoid dendritic cells (**Figure 5F, Supplementary Figure 4C**) (10).

## Discussion

This work establishes that STING has a functional role in the thymic epithelium that can impact thymocyte development and selection. We found that activation of thymic STING significantly upregulated type I interferon signaling and unexpectedly impaired autophagic flux and the processing and presentation of peptide antigens. The changes to the thymic epithelium mediated by activated STING altered thymocyte maturation and caused both a defect in negative selection and shift in the T cell repertoire. Importantly, we made these observations not only in the presence of pathogenic *COPA* mutations that cause autoimmune disease, but also after administering a systemic STING agonist being trialed for cancer patients. Thus, our data has broad implications for any setting that activates thymic STING including not only the clinical scenarios we studied, but also in patients with systemic infections or other more common autoimmune syndromes such as systemic lupus erythematosus (SLE). Furthermore, these findings are relevant throughout the life span, given the recent landmark study that showed the thymus modulates the risk of both cancer and autoimmunity in adults (37).

The major STING signaling output in thymic epithelial cells we identified was type I interferons. There has been increasing interest to define how interferons in the thymus affect developing T cells, especially after it was shown that TECs have high levels of constitutive *IFN-β* expression under homeostatic conditions (38). Further investigation is needed to understand how interferon secretion in the thymus is regulated, particularly given recent studies that suggest loss of interferon expression in the thymus may lead to the generation of interferon autoantibodies that predispose to severe COVID-19 and other infections (39). In addition to nucleic acid sensing molecules such as STING, the transcriptional activator Autoimmune regulator (AIRE) may have a role in controlling interferon expression in TECs (40). Interestingly, a number of different viruses, including severe acute respiratory syndrome coronavirus 2 (SARS-CoV-2), have been shown to directly target the thymic microenvironment and infect cells involved in thymocyte selection, including thymic epithelial cells (41, 42). Although we have not yet shown it directly, our work suggests that systemic viral or bacterial infections have the potential to activate thymic STING and alter T cell selection which could in theory trigger autoimmune disease. Indeed, investigators have shown that viruses may disrupt central tolerance and promote autoimmunity by reducing thymic epithelial cell numbers or the expression of tissue specific antigens (43). The activation of STING (and possibly other nucleic acid sensing molecules) in the thymus demonstrates another mechanism by which activation of an innate immune signaling pathway leads to loss of self-tolerance.

Our study has important implications for understanding COPA syndrome pathogenesis and potentially other disorders that cause constitutive activation of STING (44, 45). COPA syndrome was originally characterized as an autoimmune disorder given the presence of high titer autoantibodies and the significant expansion and skewing of adaptive T helper cell subsets (2). Indeed, T cells appear to have an important role in disease pathogenesis and COPA syndrome patients respond well to mycophenolate moefitil, which significantly dampens adaptive immune cell subsets (3, 4). Future studies might evaluate whether other patient populations including those with STING-associated vasculopathy with onset in infancy (SAVI) also have alterations in T cell selection (45).

There has been an intense interest to modulate STING signaling therapeutically given its central role in a broad range of biological contexts important to human health and disease (46). Our ability to activate STING in the thymus with small molecules and alter T cell selection is highly relevant for cancer treatment protocols that employ these drugs (47). We showed not only how STING agonists can enhance anti-tumor T cell responses, but also pinpointed a potential mechanism by which STING agonists might provoke autoimmune reactions. When administered with checkpoint inhibitors, STING agonists acting on the thymus may predispose to immune related adverse events (48). In any case, we demonstrate the critical importance of studying off-target effects of systemically administered STING therapeutics (49), and our discovery of STING’s functional role in the thymus will be similarly important to study in clinical contexts that trial STING inhibitors.

Our study has limitations. Persistent activation of thymic STING lead to an expected increase in type I interferons but it remains unclear why macroautophagic flux was impaired. Activation of STING has been shown to induce non-canonical autophagy (22) although this degradation pathway is not known to have a role in antigen processing and presentation. Future study is needed to understand how persistent STING activation impacts STING-induced autophagy and whether it directly affects macroautophagic function in TECs. Other questions requiring further exploration include understanding whether type III interferons are involved in mediating thymic selection in our system. An analysis of systemic STING agonists on other stromal and hematopoietic populations in the thymic environment is also needed.

We uncovered an unexpected and novel role for STING in thymic epithelial cell function and provide new insight into how STING shapes the T cell repertoire to contribute to autoimmunity and immune dysregulation. These findings have important implications for any setting in which systemic inflammation activates STING in the thymic stroma. This includes not only COPA syndrome, but potentially other autoimmune diseases such as SLE. In addition, our findings have relevance for STING agonist use in cancer or understanding how T cell responses might be altered during systemic infections that activate thymic STING.

## Acknowledgements

We thank Mark S. Anderson and Michael Waterfield for critical review of the manuscript. Han Yin and Joe Germino for assistance with data handling. Microscope funding acknowledgment: UCSF Program for Breakthrough Biomedical Research funded in part by the Sandler Foundation and the Strategic Advisory Committee and the EVCP Office Research Resource Program Institutional Matching Instrumentation Award.

## Funding

- National Institutes of Health grant R01AI168299 (CSL, SK, NS, AKS)
- National Institutes of Health grant R01AI137249 (ZD, CSL, AKS)
- Microscope funding acknowledgment: UCSF Program for Breakthrough Biomedical Research funded in part by the Sandler Foundation and the Strategic Advisory Committee and the EVCP Office Research Resource Program Institutional Matching Instrumentation Award.

## Author contributions

- Conceptualization: AKS
- Methodology: ZD, CSL, AKS
- Investigation: ZD, CSL, SK
- Visualization: ZD, CSL, SK, NS
- Funding acquisition: AKS
- Project administration: ZD, AKS
- Supervision: AKS
- Writing – original draft: ZD, AKS
- Writing – review & editing: AKS

## Competing interests

Authors declare that they have no competing interests.

## Materials and Methods

### Mice

*Copa^E241K/wt^* mice were previously generated in our laboratory (5). The B6(Cg)-Ifnar1^tm1.2Ees^/J (Ifnar1^-^), B6.Cg-Tg(TcraTcrb)425Cbn/J (OT-II), C57BL/6J-Sting1^gt^/J, C57BL/6-Tg(CAG-RFP/EGFP/Map1lc3b)1Hill/J (CAG-RFP-EGFP-LC3), B6.129S7-Rag1^tm1Mom^/J (Rag1 KO), B6.Cg-Rag1^tm1Mom^ Tyrp1^B-w^ Tg(Tcra,Tcrb)9Rest/J (RAG1^-^ B^W^ TRP-1 TCR) mice were acquired from the Jackson Laboratory. GFP-LC3 transgenic mice (25) were provided by Jayanta Debnath. All mice were housed in a specific pathogen-free facility at UCSF, and all protocols were approved by UCSF’s Institutional Animal Care and Use Committee.

### Histology and confocal microscopy

Isolated thymi were fixed for one hour in PBS with 4% PFA, cryopreserved overnight in 30% sucrose, embedded in Tissue-Tek OCT (Sakura Finetek USA), and frozen sectioned at 10 µm. Tissue sections were stained with antibodies against phosphorylated STING (D8F4W, Cell Signaling Technology), cytokeratin 5 (EP1601Y, Abcam), and cytokeratin 8 (EP1628Y, Abcam). Fluorescent conjugated secondary antibodies were acquired from Thermo Fisher Scientific, and sections were counterstained with DAPI (BioLegend). Images were captured with a CREST LFOV Spinning Disk/C2 confocal.

### Thymic epithelial cell isolation

Thymic epithelial cells were isolated as previously described (6). In brief, isolated thymi from 4-week-old mice were minced with razor blades on ice, digested three times (DMEM high glucose with 2% FBS, 100 µg/ml DNase I, and 100 µg/ml Liberase TM, Roche) or until no tissue fragments remained, and quenched and washed with magnetic activated cell sorting buffer (PBS with 5 mg/ml BSA and 2 mM EDTA). Cells were fractionated with a Percoll (Cytiva) gradient of densities 1.115, 1.065, and 1.0. Thymic epithelial cells were isolated from the interphase of 1.065 and 1.0 densities and further analyzed by flow cytometry.

### RNA sequencing and data analysis

RNA sequencing of murine thymic epithelial cells was previously generated (6). In brief, cDNA was generated from isolated RNA with the Nugen Ovation method (7102-A01, Tecan), and a sequencing library was created with the Nextera XT method (FC-131-1096, Illumina). Libraries were sequenced on Illumina HiSeq4000 and resulting reads were aligned to ensembl build GRCm38.78 with STAR (v2.4.2a). Read counts per gene were inputted to DESeq2 (v1.26.0) and resulting differential gene expression profile was plotted with EnhancedVolcano (v1.18.0).

Human *STING1* transcript expression was assessed with a publicly available cell atlas of human thymus(35) (https://doi.org/10.5281/zenodo.5500511) and Python toolkits Scanpy (v1.9.4), pandas (v2.0.3), and NumPy (v1.25.0).

### Flow cytometry and antibodies

Single cell suspensions of thymocytes and splenocytes were prepared by mechanically disrupting the thymus and spleen and passing the cells through 64 µm filters (Genesee Scientific). Following red blood cell lysis (BioLegend), cells were maintained on ice in RPMI 1640 supplemented with 5% FBS. An aliquot of each cell suspension was mixed with AccuCheck Counting Beads (PCB100, Invitrogen), analyzed by flow cytometry, and total cell number was calculated according to manufacturer’s protocol.

For evaluation of surface markers, cells were blocked with 10 µg/ml anti-CD16/32 for 15 minutes at room temperature and then stained with indicated antibodies in FACS buffer (PBS with 2% BSA) on ice for one hour.

For stimulation of intracellular cytokines, freshly isolated splenocytes were stimulated in RPMI 1640 with 5% FBS, 10 ng/ml PMA (Sigma-Aldrich), 1 µg/ml ionomycin (Sigma-Aldrich), and 5 µg/ml Brefeldin A (Biolegend) at 37°C for four hours.

For intracellular staining, cells were first stained with Zombie Violet viability dye (BioLegend), surface stained, fixed (PBS with 4% PFA), permeabilized (PBS with 0.2% saponin), and lastly stained with antibodies against intracellular targets in the presence of permeabilization buffer overnight at 4°C. Flow cytometry data were acquired on FACSVerse or LSRFortessa (BD Biosciences) and analyzed with FlowJo v10.

Antibodies to the following were purchased from BD Biosciences: mouse TCR Vβ screening panel (557004), Vβ14 (14–2); BioLegend: CD3 (145-2C11), CD5 (53-7.3), CD19 (6D5), CD25 (PC61), CD45 (30-F11), CD62L (MEL-14), CD80 (16-10A1), EpCAM (G8.8), H-2 MHC class I (M1/42), I-A^b^ MHC class II (AF6-120.1), IFNγ (XMG1.2), Ly-51 (6C3), NK1.1 (PK136), TCRγ/δ (GL3), TNFα (MP6-XT22), Vα2 (B20.1), Vβ5 (MR9-4); Cell Signaling Technology: cleaved caspase 3 (D3E9); Cytek Biosciences: B220 (RA3-6B2), CD4 (RM4-5), CD8α (53-6.7), CD11c (N418), CD44 (IM7), CD69 (H1.2F3), F4/80 (BM8.1), Gr-1 (RB6-8C5), TCRβ (H57-597); Thermo Fisher Scientific: Nur77 (12.14); and UCSF’s Hybridoma Core Facility: CD16/CD32 (2.4G2). AccuCheck Counting Beads were purchased from Thermo Fisher Scientific.

### Quantitative real-time PCR

RNA was isolated with EZNA Total RNA kit (Omega Bio-tek) and reverse transcribed to cDNA with SuperScript III and oligo d(T)_16_ primers (Thermo Fisher Scientific). Quantitative PCR was performed on a Bio-Rad CFX thermal cycler with TaqMan Gene Expression assays from Thermo Fisher Scientific: Gapdh, Mm99999915_g1; Ifit1, Mm00515153_m1; Isg15, Mm01705338_s1.

### Immunoblotting and antibodies

Cells were lysed in Cold Spring Harbor NP-40 lysis buffer (150 mM NaCl, 50 mM Tris, pH 8.0, and 1.0% Nonidet P-40) containing protease and phosphatase inhibitors (PMSF, NaF, Na_3_VO_4_, and Roche PhosSTOP). Lysates were cleared by centrifuging at 10,000 g for 10 min at 4° C, size separated on SDS-PAGE gels, and wet transferred onto polyvinylidene fluoride membrane. Membranes were blocked in tris-buffered saline with 0.05% Tween 20 (TBS-T) and 5% nonfat dry milk for 1 h at room temperature, followed by overnight incubation at 4° C with primary antibodies diluted in TBS-T with 5% BSA. Membranes were washed three times with TBS-T for 10 min, incubated with HRP-conjugated IgG secondary antibody (Jackson Immunoresearch) for 1 h at room temperature, washed three times with TBS-T for 10 min followed once with TBS for 10 min. Lastly bands were visualized with SuperSignal West Femto Chemiluminescent Substrate (Thermo Fisher Scientific) and Bio-Rad’s ChemiDoc MP imager. Antibodies to the following were purchased from Cell Signaling Technology: LC3B (D11), Phospho-STING (D8F4W); and Santa Cruz Biotechnology: GAPDH (6C5).

### Plasmids and transient transfection

FLAG-tagged wild type and E241K human COPA expression plasmids were previously described(2). EGFP-tagged human STING lentiviral plasmid was a gift from Dr. Tomohiko Taguchi (Tohoku University, Sendai, Japan). HEK293T cells (ATCC) were seeded at 40-60% density, transiently transfected with jetOPTIMUS reagent (Polyplus), and lysed 48 hours later for immunoblotting.

### Bone marrow chimeras

Host mice aged 3-4 months old were fasted overnight and then depleted of bone marrow with two doses (550 rads each, separated 12 h apart) of γ radiation. Donor bone marrow cells were isolated from femurs and tibias of 8 weeks old, gender-matched mice. T cells were removed with CD90.2 antibody (30-H112, BioLegend) and Dynabeads (65601, Thermo Fisher Scientific). Ten million bone marrow cells suspended in 200 µl of saline were intravenously transferred into each host. Chimeras were housed under specific pathogen free conditions and analyzed 6 to 8 weeks after reconstitution.

### In vivo STING activation

Beginning at 5-7 weeks of age, STING agonist diABZI compound 3 (ProbeChem) was administered intraperitoneally (0.25 mg/kg) to *Rag1^-/-^ Tyrp1^B-w/wt^* TRP-1 TCR mice every other day, seven times, followed by seven days of rest. Mice were then euthanized to harvest cells from thymi and peripheral lymphoid organs for analysis or adoptive transfer.

### Tumor challenge

*Copa^E241K/wt^* and wild type littermates were subcutaneously challenged with 1 × 10^5^ B16-F10 melanocytes (a gift from Dr. Lawrence Fong, UCSF) at the left flank. Tumor volume was estimated as length × width^2^/2, and a volume of 1,000 mm^3^ was classified as death for Kaplan-Meier survival analysis.

For adoptive transfers, *Rag1^-/-^* mice were subcutaneously challenged with 5 × 10^4^ B16-F10 cells at the left flank on day 0. Mice received 500 rads of X-ray radiation (X-Rad320, Precision X-Ray Irradiation) and 125 µg of PD-1 antibody (CD279, BioXCell) i.p. on day 2. Total splenocytes from one *Rag1^-/-^ Tyrp1^B-w/wt^* TCR mice given STING agonist or vehicle, as described above, were transferred into one tumor challenged Rag1^-/-^ mice on also on day 2. Mice received additional 125 µg of PD-1 antibody i.p. every four days. Mice were euthanized sixteen to seventeen days following tumor inoculations.

### Statistical analysis

Data is presented relative to the mean of wild type or agonist treated. Statistical analysis was performed with Prism 9 (GraphPad) or R 4.0.0 (R Foundation for Statistical Computing). Unpaired, parametric, two-tailed Student’s t-test or two-tailed Mann-Whitney U test was used where indicated. Log-rank test was used for Kaplan-Meier survival analysis. P < 0.05 was considered statistically significant.

## Notes

### Competing Interest Statement

The authors have declared no competing interest.

