## Supplementary Data for "Activated STING in the thymus alters T cell development and selection leading to autoimmunity"

### Supplementary Fig 1

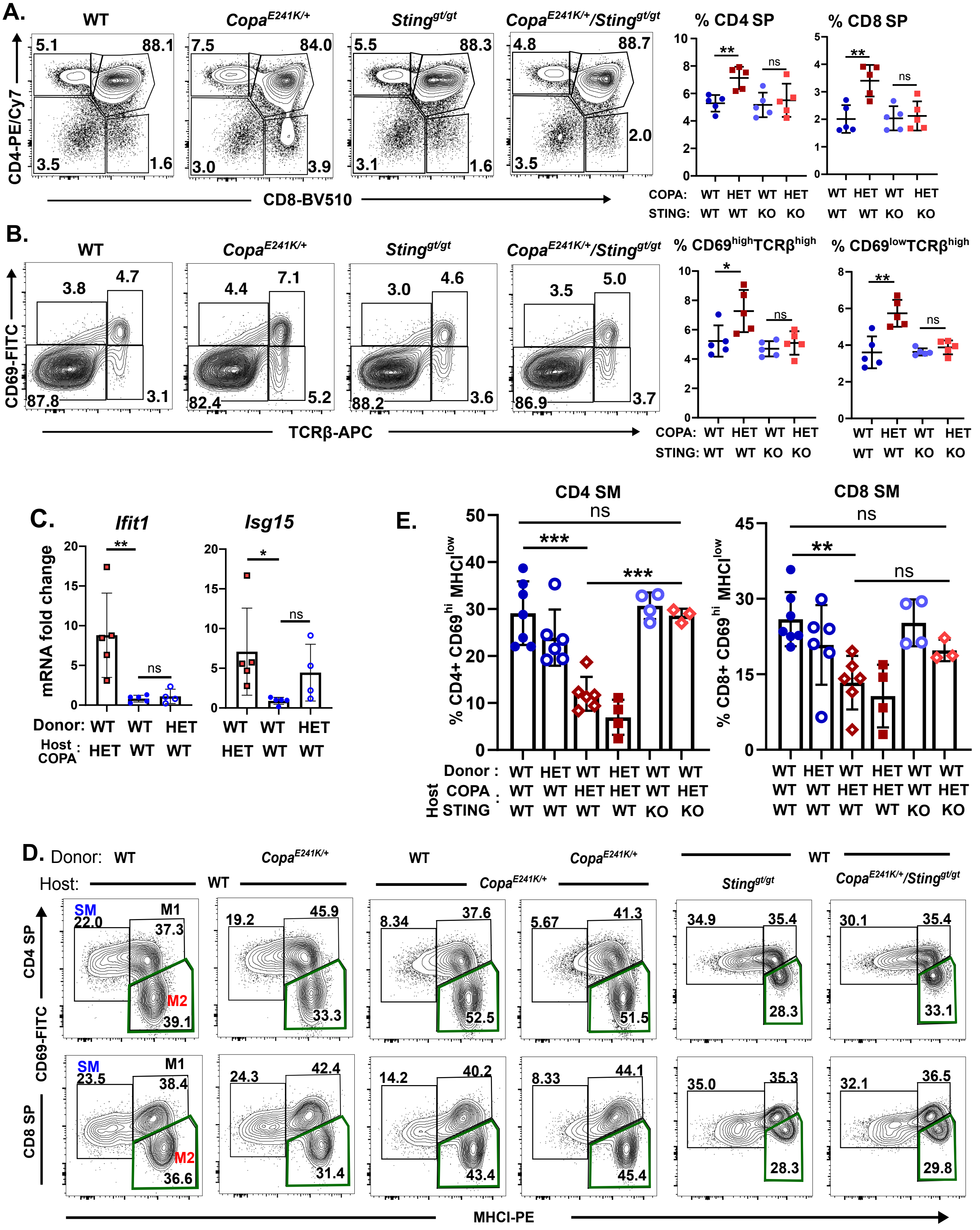

**Fig S1. (A)** Left: CD4 and CD8 profile of thymocytes. Right: percentages of single positive CD4 and CD8 thymocytes in indicated mice ($n$ = 5 for all genotypes). (**B**) Left: thymocytes expression of CD69 and TCR beta chain. Right: percentage of $CD69^{high}$ $TCR\beta^{high}$ and $CD69^{low}$ $TCR\beta^{high}$ thymocytes in indicated mice ($n$ = 5 for all genotypes). (**C**) Relative expression of interferon stimulated genes *Ifit1* and *Isg15* transcript in thymic stroma of indicated chimeras (WT→$Copa^{E241K/+}$, $n$ = 5; WT→WT, $n$ = 5; $Copa^{E241K/+}$→WT, $n$ = 4). (**D**) Flow analysis of CD69 versus MHC-I on reconstituted single positive thymocytes in bone marrow chimeras indicating SM, M1 and M2 populations in CD4 and CD8 cells. (WT→WT, $n$ = 7; WT→$Copa^{E241K/+}$, $n$ = 6; $Copa^{E241K/+}$→WT, $n$ = 6; $Copa^{E241K/+}$→$Copa^{E241K/+}$, $n$ = 4; WT→$Sting^{gt/gt}$, $n$ = 4; WT→$Copa^{E241K/+}$ × $Sting^{gt/gt}$, $n$ = 3) (**E**) Percentages of $CD69^{high}MHCI^{low}$ (semi-mature: SM) cells among the reconstituted CD4 and CD8 single positive thymocytes in BM chimeras. Data are mean ± SD. Unpaired, parametric, two-tailed Student's *t*-test was used for statistical analysis. $p < 0.05$ is considered statistically significant. ns: not significant.

#### Supplementary Fig 2.

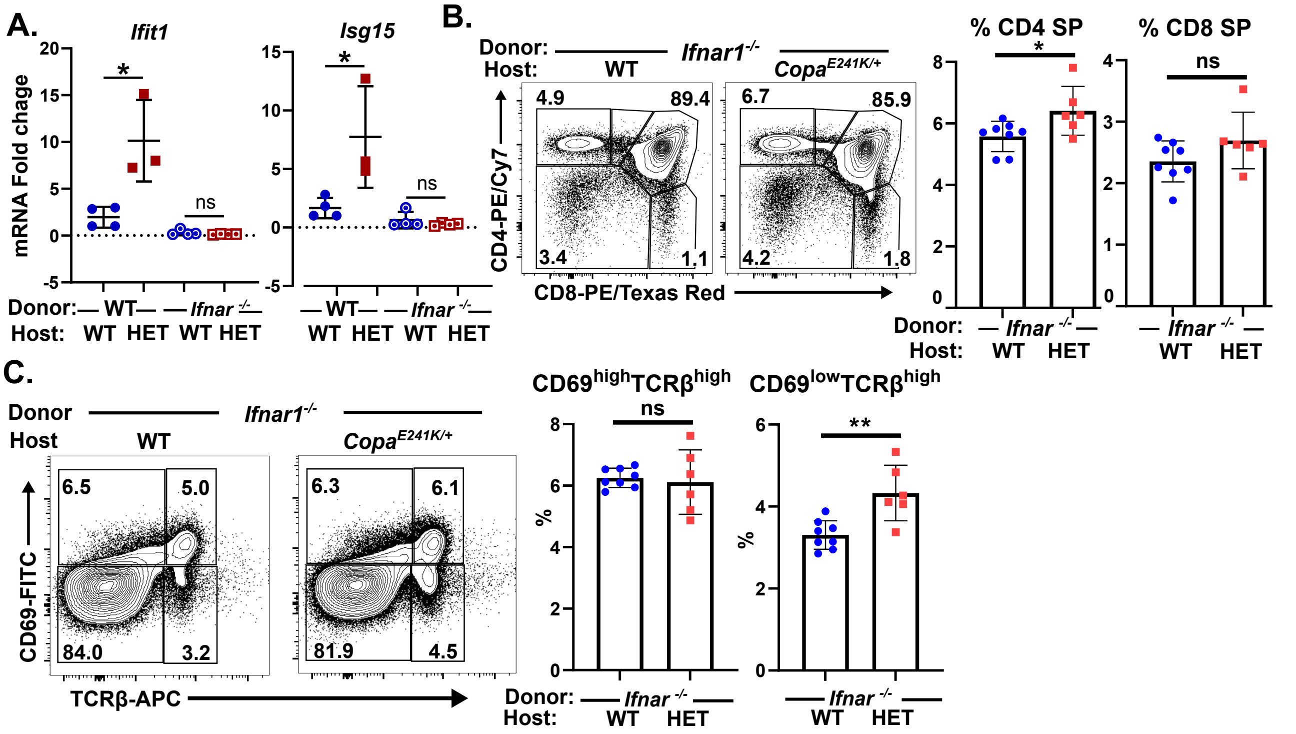


**Fig S2. (A)** Relative expression of interferon stimulated genes *Ifit1* and *Isg15* transcript in thymic stroma of indicated chimeras (WT→WT, n = 4; WT→*Copa*$^{E241K/+}$, n = 4; *Ifnar1*$^{-/-}$→WT, n = 4; *Ifnar1*$^{-/-}$→*Copa*$^{E241K/+}$, n = 4). **(B)** Left: CD4 and CD8 profile of thymocytes in *Ifnar1*$^{-/-}$ chimeras. Right: percentage of single positive CD4 and CD8 thymocytes in indicated chimeras (*Ifnar1*$^{-/-}$→WT, n = 7; *Ifnar1*$^{-/-}$→ *Copa*$^{E241K/+}$, n = 5). **(C)** Left: CD69 and TCR beta chain profiling in reconstituted thymocytes of *Ifnar1*$^{-/-}$ chimeras. Right: percentages of CD69$^{high}$ TCRβ$^{high}$ and CD69$^{low}$ TCRβ$^{high}$ thymocytes in indicated chimeras (*Ifnar1*$^{-/-}$→WT, n = 7; *Ifnar1*$^{-/-}$→ *Copa*$^{E241K/+}$, n = 5). Data are mean ± SD. Unpaired, parametric, two-tailed Student's t-test was used for statistical analysis. $p < 0.05$ is considered statistically significant. ns: not significant.

#### Supplementary Fig 3.

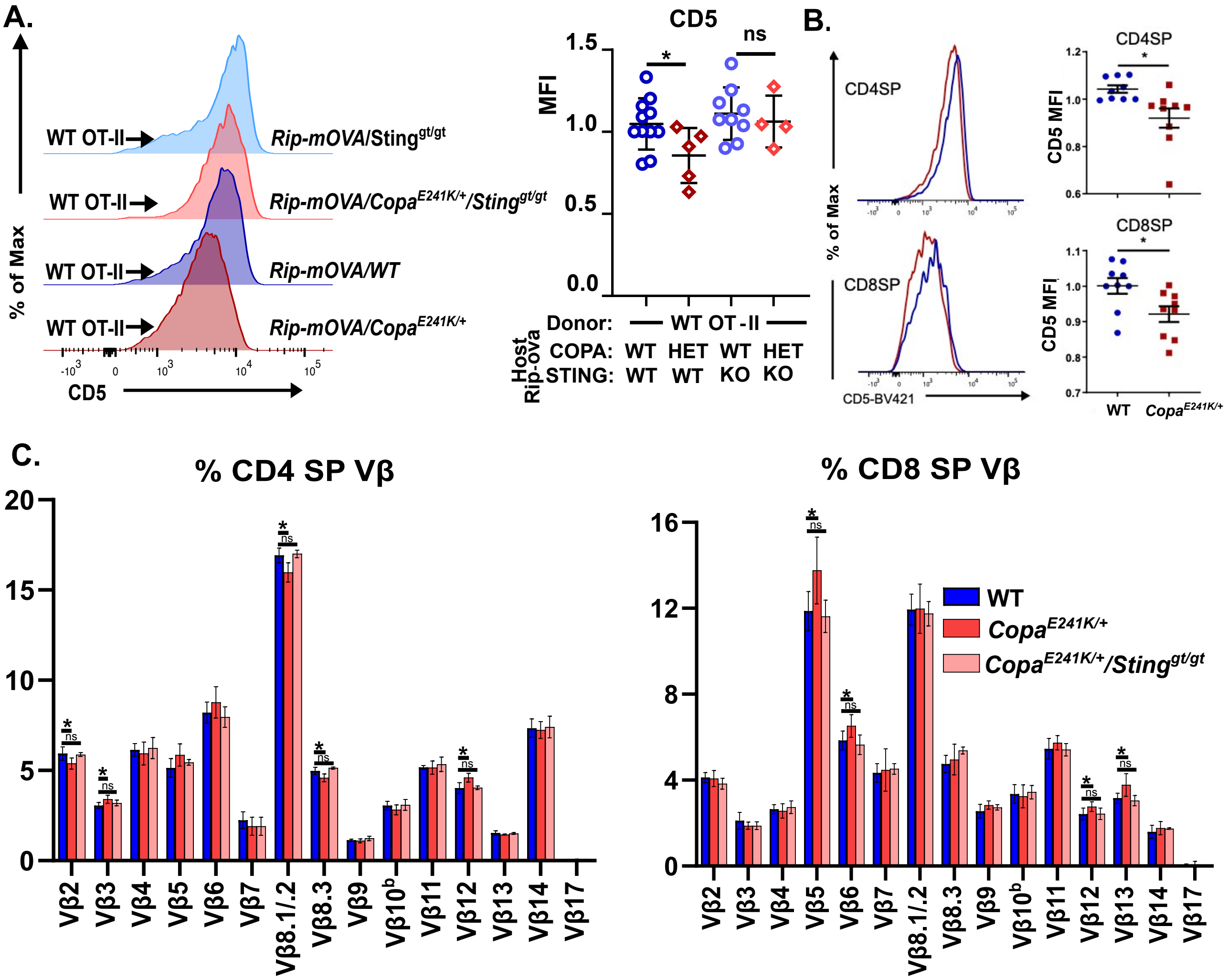


**Fig S3. (A)** CD5 expression in *Rip-mOVA/Copa/Sting* chimeras that received OT-II bone marrow. Left: representative histogram of CD5 expression; and right: CD5 MFI for indicated chimeras (OT-II→WT, n = 11; OT-II→*Copa*$^{E241K/+}$, n = 5; OT-II→*Sting*$^{gt/gt}$, n = 9; OT-II→ *Copa*$^{E241K/+}$ × *Sting*$^{gt/gt}$, n = 4) **(B)** CD5 expression in single positive thymocytes of *Copa*$^{E241K/+}$ mice. Left: representative histogram of CD5 levels; and right: MFI for indicated mice (WT, n = 9; *Copa*$^{E241K/+}$, n = 9). **(C)** Quantitation of TCR Vβ repertoire of CD4 and CD8 single positive thymocytes. (WT, n = 5; *Copa*$^{E241K/+}$, n = 4; *Copa*$^{E241K/+}$ × *Sting*$^{gt/gt}$, n = 5). Unpaired, parametric, two-tailed Student's t-test was used for statistical analysis. $p < 0.05$ is considered statistically significant. ns: not significant.

#### Supplementary Fig 4.

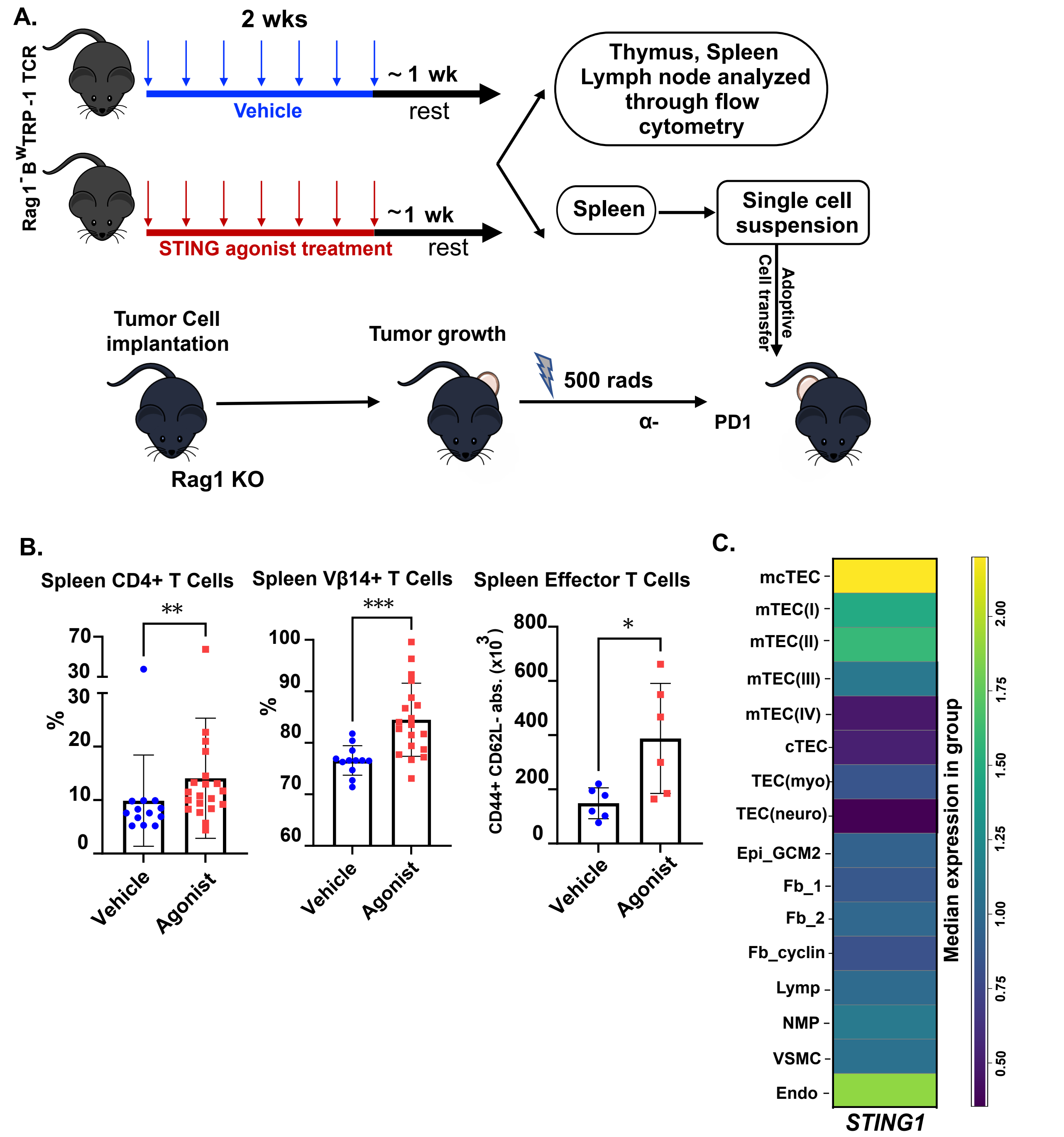


**Fig S4. (A)** Schematic for administering vehicle or diABZI STING agonist to *Rag1*$^{-/-}$ *Tyrp1*$^{B-w/wt}$ TCR mice and B16 tumor cell inoculation with adoptive cell transfer into *Rag1*$^{-/-}$ mice. **(B)** Left: splenic percentage CD4 single positive thymocytes; and middle: percentage V$\beta$14$^{+}$ CD4 SP thymocytes in vehicle and STING agonist treated *Rag1*$^{-/-}$ *Tyrp1*$^{B-w/wt}$ TCR mice (vehicle n = 14, agonist n = 21). Right: absolute number of effector memory cells (vehicle n = 6, agonist n = 6).**(C)** Matrix plot of median transcript expression of *STING1* in human thymic stromal cells. Two-tailed Mann-Whitney U-test was used for statistical analysis. $p < 0.05$ is considered statistically significant.
